# Quantifying heterogeneity of stochastic gene expression

**DOI:** 10.1101/316166

**Authors:** Keita Iida, Nobuaki Obata, Yoshitaka Kimura

## Abstract

The heterogeneity of stochastic gene expression, which refers to the temporal fluctuation of the gene product and its cell-to-cell variation, has attracted considerable interest from biologists, physicists, and mathematicians. The dynamics of protein production and degradation have been modeled as random processes with transition probabilities. However, there is a gap between theory and phenomena, particularly in terms of analytical formulation and parameter estimation. In this study, we propose a theoretical framework in which we present a basic model of a gene regulatory system, derive a steady-state solution, and provide a Bayesian approach for estimating the model parameters from single-cell experimental data. The proposed framework is demonstrated to be applicable for various scales of single-cell experiments at both the mRNA and protein levels, and it is useful for comparing kinetic parameters across species, genomes, and cell strains.

## 1. Introduction

Gene expression in prokaryotes and eukaryotes is exposed to various molecular noises. Modern single-cell gene expression analyses have revealed that gene expression levels fluctuate in each cell [1, 2, 3] and differ from cell to cell even within a clonal population in an identical environment [4, 5, 6, 7]. Meanwhile, ordered dynamics—ranging from microscopic to macroscopic levels, such as nearly perfect DNA replications [8], cell polarization [9], and mammalian embryogenesis [10]—have also been observed. These discoveries reveal that cells integrate both noisy and accurate molecular processes to make them well organized overall, demonstrating the difference between organisms and machinery. However, the question of how do cells achieve orchestration through stochastic expression kinetics arises.

It has been proposed that stochastic expression kinetics is related to its cell-to-cell variation [5], cellular memory [11], cell differentiation [12, 13], and evolution [14]. Accordingly, heterogeneous cellular responses to environmental changes have been thoroughly studied. For example, the *lac* operon in *Escherichia coli* [15, 16, 17] and *GAL* genes in *Saccharomyces cerevisiae* [18, 19, 20] produce discriminative unimodal and bimodal distributions of the protein concentration. Another study on population survival suggested that increasing expression noise, rather than the mean expression level, could provide cells with a selective advantage under stress conditions [21, 22]. These observations indicate that cells utilize the protein distribution to adapt to environmental changes. Nevertheless, many previous studies have focused on the mean and variance, not on the benefits gained from the distribution.

To address the aforementioned issue, theoreticians have developed analytical procedures to derive mRNA and protein distributions [23, 24, 25, 26, 27, 28, 29, 30, 31, 32, 33, 34]. The dynamics of protein production and degradation have been modeled by a discrete stochastic model, whereas the dynamics of transition probability have been modeled by a master equation. However, the connection between stochastic processes and probability distribution is less studied. This point indicates a gap between theory and phenomena because the master equation does not always represent the stochastic dynamics. Moreover, in practice, many previous studies are based on a discrete model that counts mRNA and protein copy numbers because they are discrete in nature. Meanwhile, in most experiments, the single-cell gene expressions are indirectly observed through the measurement of fluorescence intensity, thus indicating that we need a continuous model.

In the present article, we begin with the biophysical modeling of a simple gene regulatory system. First, we formulate the stochastic process of protein production and degradation coupled with an active-or-inactive genetic switch. Second, we introduce a system of master equations and derive an important steady-state solution expressing protein distribution. We show that the solution can be fit to experimental data with an arbitrary measurement scale. Finally, we apply the proposed theory to the thiomethylgalactoside (TMG)-induced system of *lacZYA* expression in *E. coli*, estimate the model parameters from published experimental data [15] with Markov chain Monte Carlo (MCMC) methods, and investigate the heterogeneous responses of the *lac* genes to extracellular TMG concentrations. Consequently, the results of this study demonstrate that the proposed theoretical framework is widely applicable to various types of singlecell experiments at both the mRNA and protein levels, such as reporter assays [35, 36, 37], MS2-GFP system [2, 3], fluorescent in situ hybridization [7, 36], and flow cytometry [20, 22].

## 2. Methods

### 2.1 Model

Starting with a simplified gene regulatory system, as shown in Fig. 1(a), we assume the following three phases: phase I, i.e., the extracellular phase, is a molecular reservoir in which concentrations remain constant; phase II, i.e., the cytoplasmic phase, contains a vast amount of protein molecules, and the system size is large enough to keep the condition dilute and well stirred (to benefit from the deterministic chemical reaction rate equation [38]); and phase III, i.e., the gene-mRNA phase, is a small-size subsystem in which non-negligible molecular noises exist. We further assume that the system consists of three variables, *x, y*, and *z*, and a gene-protein interaction, *f,* where *x* = 1 and *x* = 0 denote the active and inactive states of the gene, respectively, and *y* and *z* are the amounts of the synthesized protein Y and a certain effector protein Z, respectively. For the gene-protein interaction, we assume that the probability of *x* switching between 0 and 1 per unit time d*t* is described by *x* + *f* (*x*)d*t*, where

**Figure 1:**
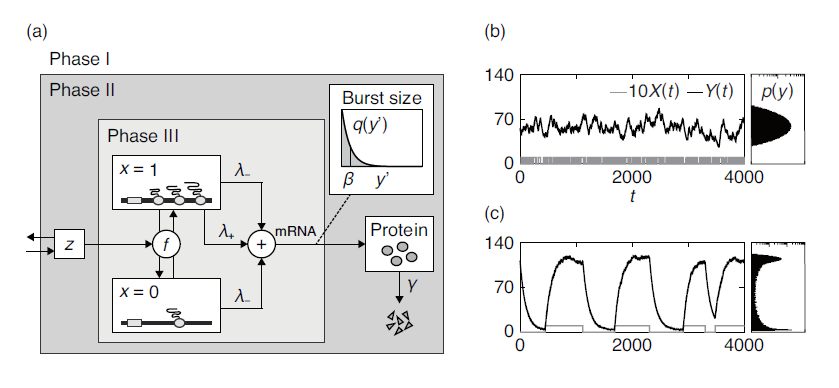
(a) Schematic representation of the simplified gene regulatory system. (b) and (c) (left) Typical sample paths of 10*X*(*t*) (gray line) and *Y* (*t*) (black line), and (right) *p*(*y*), i.e., the probability density of *Y* (*t*), plotted in log10 scale. The common parameters are biologically relevant: *λ*_*-*_ = 0.5 (this is slightly larger than the presumable value), *λ*_+_ = 15, *β* = 0.074, and *γ* = 0.01; and the control parameters (b) *k*_on_ = *k*_off_ = 10*γ*, and (c) *k*_on_ = *k*_off_ = *γ/*10.

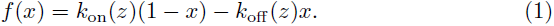

Here, *k*_on_(*z*) and *k*_off_ (*z*) are the transition rates. Assume that the influx and efflux of Z through the cytoplasmic membrane are always in equilibrium [39, 17], and let *z* always be constant. Henceforth, let *k*_on_(*z*) = *k*_on_, and let *k*_off_ (*z*) = *k*_off_.

Based on the concept of fast-slow dynamics [27, 40], we assume that the unit time is considerably larger and smaller than the lifetimes of the mRNA and protein, respectively, and Ys are produced through a bursting manner from their mRNA in the genetic phase, whereas they are degraded in a continuous manner in the cytoplasmic phase. We also define the following: *λ*_*-*_ and *λ*_+_ are the burst frequencies, *β* is the mean burst size per burst, and *γ* is the decay constant. Here, *λ*_*-*_ represents the basal transcription rate from the inactive promoter, which is referred to as “promoter leakage” [34]. Based on these assumptions, we formulate the time evolutions of *X ∈* {0, 1} and *Y ∈* R_*>*0_ by the following mixed random process:

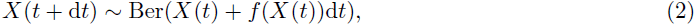

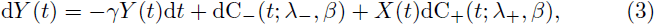

where “*∼*” denotes random sampling, and Ber(*p*) is the Bernoulli distribution, whose random variable takes a value of 1 with the probability of *p*; otherwise, it is 0. Here, dC_*i*_(*t*) (*i ∈* {+, *-*}) is the compound Poisson white noise [41, 42] defined by

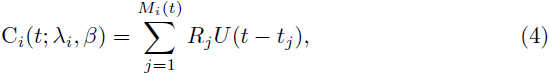

where {*M*_*i*_(*t*)} denotes a homogeneous Poisson counting process with the occurrence rate *λ*_*i*_, *U* (*t*) is the unit step function, *t*_*j*_ is the *j*th arrival time, and {*R*_*j*_} is a sequence of independent identically distributed random burst sizes with a mean of *β*. Based on experimental observations [43] and theoretical assumptions [24, 26, 27], we assume that *R*_*j*_ follows an exponential distribution with the assigned probability density 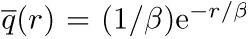. The typical sample paths of *X*(*t*) and *Y* (*t*) for different values of *k*_on_ and *k*_off_ are shown in Figs. 1(b) and (c). Note that directly predicting the values of *k*_on_ and *k*_off_ may be possible from Fig. 1(c), but such a prediction is difficult from Fig. 1(b). Hence, we need a theoretical procedure to estimate these parameters in any case.

### 2.2. Analysis

Next, we investigate the probability density functions (PDFs) of *X*(*t*) and *Y* (*t*). Since dC_*-*_(*t*) and dC_+_(*t*) are independent random processes, we can analyze the following systems individually:

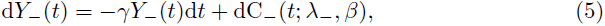

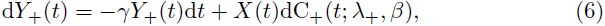

where *Y* (*t*) = *Y*_*-*_(*t*)+*Y*_+_(*t*). Let *p*(*t, y*), *p*_*-*_(*t, y*), and *p*_+_(*t, x, y*) be the transition PDFs of *Y* (*t*), *Y*_*-*_(*t*), and *Y*_+_(*t*), respectively, and let 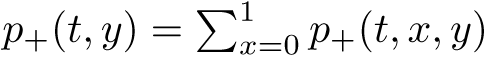. According to the procedure in references [41, 42], we obtain the following system of integro-differential equations along with the normalization condition:

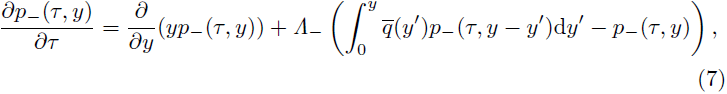

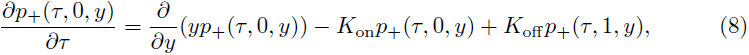

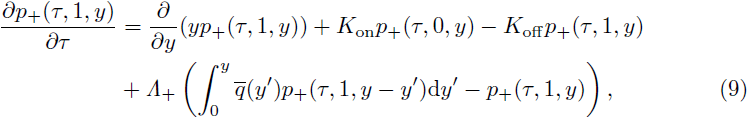

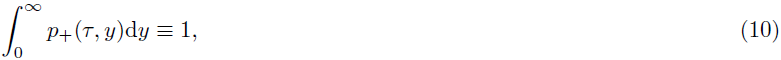

where *τ* = *γt, K*_on_ = *k*_on_*/γ, K*_off_ = *k*_off_ */γ*, and *Λ*_*i*_ = *λ*_*i*_*/γ* (*i ∈* {+, *-*}). Note that Eq. (7) is the so-called Kolmogorov-Feller equation [44] introduced by Friedman and his coworkers as a model of constitutive gene expression [26].

Let us consider the steady-state problem by setting the left-hand sides of Eqs. (7)–(9) to zero. Let 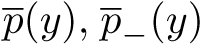 and 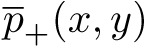 be the PDFs under the steady state, and let 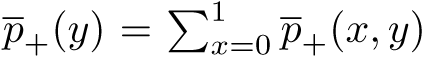. The Laplace transforms of Eqs. (7)–(9) yields

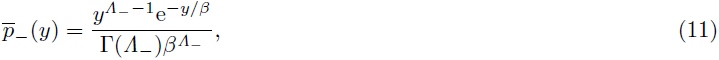

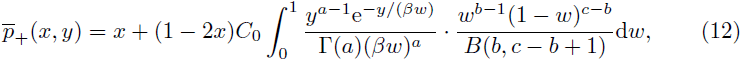

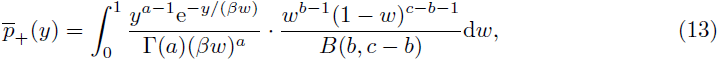

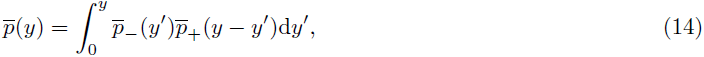

where *C*_0_ = *K*_off_ */*(*K*_on_ + *K*_off_), *a* + *b* = *K*_on_ + *K*_off_ + *λ*_+_, *ab* = *K*_on_*λ*_+_, and *c* = *K*_on_+*K*_off_ (see the Supplemental Material for the derivation). Here, Eq. (13) is the PDF of the weighted gamma distribution, whose scale parameter *βw* is averaged over *w ∈* (0, 1) with the assigned beta distribution as a weight function.

Assuming that *K*_on_, *K*_off_ *≪* 1, Eq. (13) can be approximated as follows:

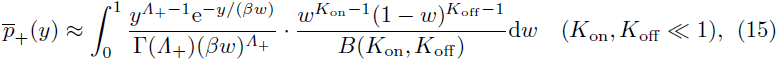

where *Λ*_+_, *K*_on_, and *K*_off_ serve as the shape parameters of the protein distribution, and *β* is the scale parameter. In fact, as shown in Eqs. (11)–(13), changing the value of *β* only results in the scaling of 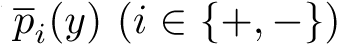. In other words, for any *k >* 0, the following identity holds:

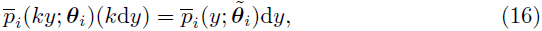

where ***θ***_*i*_ = (*k*_on_, *k*_off_, *λ*_*i*_, *β, γ*) and 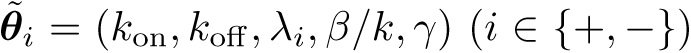. This result indicates that Eqs. (11)–(14) can be fit to various experimental data with an arbitrary measurement scale of *y*. In addition, we found that the protein distribution defined by Eq. (13) is analogous to the “Poisson-Beta distribution”, which is the solution of the discrete model for the birth-and-death process of the mRNA copy number with an all-or-none genetic switch [45, 46]. Through the analysis, we can also calculate any order of moment (see the Supplemental Material for the calculation).

## 3. Results

Finally, we apply our theory to the TMG-induced system of *lacZYA* expression in *E. coli*, as reported in 2004 [15]. In the study, cells with well-defined initial states, either uninduced or fully induced, were used, and the fluorescence intensity of the green fluorescent protein (GFP) in each cell toward the various TMG concentrations was measured. The remaining definitions and assumptions in this study are as follows: (i) *x, y*, and *z* denote the activity of the *lac* promoter and the intracellular levels of the GFP and TMG, respectively; (ii) *z* is constant for the system, which is assumed to be in a steady state [15]; (iii) *p*(*y*) denotes the cell-to-cell variation of *y* at the steady state; (iv) *k*_on_ *≫ k*_off_ is defined with respect to the fully induced population; (v) *λ*_*-*_ = 0 by assuming that the genetic switch is all-or-none; (vi) *λ*_*-*_ +*λ*_+_, *β*, and *γ* are the maximum transcription rate constant defined for the fully induced population, the average scaled number of protein molecules being produced during the *lac* mRNA lifetime, and the decay constant of GFP, respectively; and (vii) (*k*_on_, *k*_off_) and (*λ*_+_, *β*) are dependent on and independent of the extracellular TMG, respectively. Based on assumptions (i)–(vii), we estimate *λ*_+_ and *β* from the experimental data with the fully induced population [15, 43, 47, 48, 49]. Here, the estimated values are *λ*_+_ = 15 min^*-*1^, *β* = 0.074, and *γ* = 0.01 min^*-*1^ (see the Supplemental Material for the parameter estimation).

As shown in the previous study, most of the GFP distributions are far from Gaussian, which indicates that such distributions are poorly characterized by the mean and variance. Hence, we adopt a Bayesian approach utilizing the Metropolis algorithm [50], and we estimate the posterior distributions of *k*_on_ and *k*_off_ from the published single-cell data [15]. Note that each posterior had a single peak over our searched parameter range. Figs. 2(a) and (b) show the estimated mean values of *k*_on_ and *k*_off_ as functions of the extracellular TMG levels, and Fig. 2(c) shows the values of 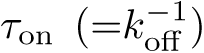 and 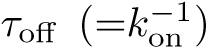, which are the average durations of *X*(*t*) = 1 and *X*(*t*) = 0, respectively, for both preuninduced and preinduced populations. Consequently, we found that the preuninduced and preinduced populations mainly modulate *k*_off_ for 3–21 and 3–5 μM TMG, respectively, the values of which are necessary for their half-induction, i.e., the bimodal distribution (see the shaded region in Fig. 2(c)). This result is comparable with those found at the mRNA level [47]. In addition, we found that the preuninduced and preinduced cells mainly modulate *k*_on_ for 24–30 and 6–30 μM TMG, respectively, the values of which are necessary for their further induction.

**Figure 2:**
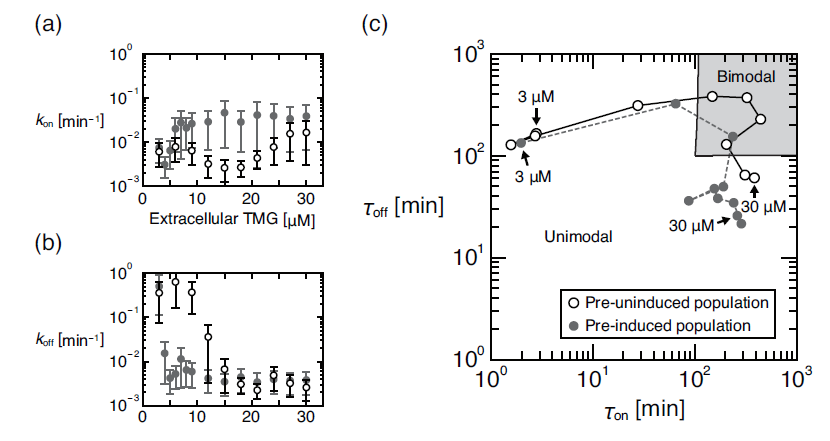
The estimated mean values of our model parameters *k*_on_ and *k*_off_ for the preuninduced (unfilled circles) and preinduced (filled circles) populations toward the various extracellular TMG concentrations. (a) and (b) The rate parameters *k*_on_ and *k*_off_ (mean *±* standard deviation) versus TMG levels. (c) The average durations of *X*(*t*) = 1 and *X*(*t*) = 0 defined by 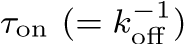 and 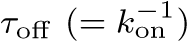, respectively, for the different TMG concentrations: 3–30 μM with 3 increments for unfilled circles, and 3–9 with 1 increment having 12, 15, and 30 μM for filled circles. The shaded region at the right top corner indicates the bimodality of 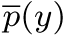. The solid and dashed lines connecting the points are simply visual guides.

## 4. Discussion

We have shown that our theoretical framework can estimate the biological parameters from single-cell data measured as fluorescence intensity. Fig. 2(c) shows that *τ*_on_ *≈* 2.78 and *τ*_off_ *≈* 164.82 min at 3 μM TMG for the preuninduced population. The experimental study with direct measurement using another *E. coli* strain under the control of a repressed *lac* promoter reported that gene expression burst lasts *∼*3 to 15 min and that the average time between two adjacent expressions is 46 min [43]. Hence, careful tuning of the parameters *λ*_*i*_ (*i ∈* {+, *-*}), *β*, and *γ* in accordance with given experimental data may help researchers accurately quantify the kinetic parameters *k*_on_ and *k*_off_.

Interestingly, Figs. 2(a) and (b) show that *k*_on_(*z*) and *k*_off_ (*z*) are nonlinear functions of TMG levels. In this case, if *z* is a random variable that obeys a distribution with nonzero variance, then the mean output value of *y* may deviate from that predicted from deterministic reaction kinetics. This situation occurs at any time when considering a random molecular flux across the cell membrane, a genetic circuit with feedback loops, and so forth. Mathematically, Jensen’s inequality may help us predict such derivation provided that *k*_on_(*z*) and *k*_off_ (*z*) are convex functions. However, Figs. 2(a) and (b) indicate that *k*_on_ and *k*_off_ are nonconvex, which leads to a notorious mixed convex problem [51].

Within the next decade, single-cell experiments with quantitative mathematical biology will enable comparing biologically important parameters such as *τ*_on_ and *τ*_off_ across species, genomes, and strains as functions of various environmental conditions, part of which were briefly reviewed by Lionnet and Singer [52]. Gnugge et al. reported that *E. coli* and *S. cerevisiae* have a similar core network in the lactose and galactose utilizing systems, respectively [53], but the functional role of the network complexity has yet to be determined. Our results show that the transition behaviors of the system can be compared on the (*τ*_on_, *τ*_off_)-plane along with the probability distributions (Fig.2(c)), which may provide a better understanding of the quantitative differences across species. Dar, Razooky et al. proposed an ingenious method for estimating their model parameters from three statistics (i.e., the coefficient of variation, expression level, and autocorrelation time) and mapped the kinetic features of human gene expression into their parameter space through a genome-wide experiment [54]. However, their method is based on a finite number of statistics, which can estimate a smaller number of parameters, and the experiment requires a high-resolution real-time monitoring, which is costly. To overcome these issues, a Bayesian or maximum likelihood approach is suitable, as Shahrezaei et al. mentioned in their report [27]. In this respect, our framework including MCMC performed well without any concern for the number of statistics. Choi, Cai et al. experimentally examined various *E. coli* strains with different genetic constructions and found that the binding affinity of the effector protein to its target loci significantly changes the protein distribution [17]. Linking DNA structure, such as looping, chemical modification, and DNA-protein complex formation, to its expression pattern is an important future work.

## 5. Conclusion

We have proposed a fairly general model of a gene regulatory system along with the clear biophysical assumptions, and we formulated it by both stochastic differential equations (Eqs. (2) and (3)) and the corresponding master equations (Eqs. (7)–(10)). We subsequently derived the steady-state solution while avoiding complicated forms such as intricate combinations of hypergeometric functions, which allows one to understand the parameters that affect the shape and scale of the protein distribution (Eq. (15)). We also demonstrated that the solution can be fit to experimental data with an arbitrary measurement scale.

As an application, we investigated the TMG-induced system of *lacZYA* expression in *E. coli*. Accordingly, we found that the system mainly modulates *k*_off_ for the lower or intermediate levels of induction and *k*_on_ for the higher level,which can be predicted on the (*τ*_on_, *τ*_off_)-plane (Fig. 2(c)). Finally, we conclude that our theoretical framework is widely applicable to various types of single-cell experiments at both the mRNA and protein levels, and it is expected to be useful for predicting the kinetic behaviors of a given biological network, including genetic oscillations [1, 55], cell differentiations [13], and evolution [22].

## Acknowledgments

The authors thank Satoshi Mizuno, Soichi Ogishima, Naoko Kasahara, Satoshi Nagaie, and Rie Norita for their helpful discussions. This work was partially supported by JSPS KAKENHI Grant Number 16H03939.

